# Fast activated outward cation channel activation allows H^+^-ATPase and anion channel reactivation upon strong depolarization in young root hairs of *Arabidopsis thaliana*

**DOI:** 10.1101/2025.03.14.643228

**Authors:** François Bouteau, David Reboutier, Aurélien Dauphin

## Abstract

Root hairs are a primary site for nutrient absorption and sensors of the root environment where signaling processes are initiated. Regulation of plasma membrane transports are fundamental for root hair tip growth, plant nutrition and responses to environmental signals. If efforts have been made to undercover the repertoires of root hair ion system transports, the roles of some of these ion transport systems remain elusive. Here we describe, using discontinuous single electrode voltage clamp, a technic that allows working in intact root hairs, the putative role of an outward current presenting a quite chaotic activation, in the polarization regulation of root hairs of *Arabidopsis thaliana*.

## Introduction

Roots hairs are tubular, single-cells that develop on the surface of roots. They emerge from the epidermal cell layer perpendicularly to the root surface. They serve to anchor the plant root system to the substrate, to increase the absorbing surface of the root, playing a major role in the uptake of nutrients and water, and finally, to interact with the rhizosheath. They behave like sensors able to perceive signals from their environment (Long, 1996; Peterson and Farquhar, 1996, Dolan, 2017; Zhang et al., 2018).

Ionic exchanges across the root hair cell plasma membrane have been shown to be fundamental for their growth, for plant nutrition and for plant microbe interactions. Root hair shape depends on a tip growth that occurs through a highest Ca^2+^ concentration close to the tip apex (Tian 2020). The Ca^2+^ gradient necessary for their growth is thought to be generated by hyperpolarization-activated calcium channels, which are localized to the plasma membrane at the hair tip. Cytoplasmic Ca^2+^ dynamics is concomitantly strongly coordinated with reactive oxygen species and apoplast pH. (Stéger Palmgren 2022). Furthermore, other channels import osmotically active K^+^ and Cl^-^ ions, which help to sustain turgor pressure as the hair grows. Beside this, Ca^2+^, Cl^-^, K^+^ and H^+^ fluxes represent one of the earliest responses to symbiotic factors (Felle et al., 1998, Kurkdjian 2000, Reboutier 2002) or pathogenic factors (Bouizgarne et al. 2004, 2006).

Using different electrophysiological techniques and molecular approaches different conductances were reported in various species. Hyperpolarization-activated and depolarization-activated calcium channels were reported at the tip of *Arabidopsis thaliana* root hairs (Very and Davies, 2000). Tian et al. (2020) reviewed recent studies that have identified cyclic nucleotide-gated channel (CNGC) members as central regulators of Ca^2+^ oscillations in the root hair tip. These include CNGC5, CNGC6, CNGC9 and CNGC14, which fulfil partially redundant functions and are essential for sustaining polarized growth of root hairs. As a major osmotically active ion, K^+^ is also expected to be assimilated to maintain cell turgor during root hair tip growth. An inwardly rectifying shaker-like conductance was detected by discontinuous single electrode voltage-clamp technique (dSEVC) in intact root hairs of *Medicago sativa*, (Bouteau et al., 1999; Kurkdjian et al., 2000) and *A. thaliana* (Dauphin et al. 2001). By using the patch-clamp technique on protoplasts Gassmann and Schroeder (1994) also described an inwardly rectifying K^+^ conductance in root hairs of wheat. *A. thaliana* and *Medicago trunculata* root hair protoplasts were shown to be equipped with two K^+^ inwardly rectifying shaker channel subunits (AKT1 and AtKC1) (Ivashikina et al., 2001; Reintanz et al., 2002, Wang et al. 2019). The involvement and AtKC1 and AtAKT1 but also of AtKUP4 (TRH1) have been recently confirmed to participate in K^+^ acquisition by using a fluorescent K^+^ sensor (Sun et al. 2024). In *M. sativa* beside this shaker-like inwardly rectifying K^+^ currents, a strongly outwardly rectifying shaker-like conductance was also evidenced by dSEVC on intact root hairs (Bouteau et al., 1999). In root hairs of *A. thaliana* such K^+^ outwardly rectifying current was attributed to Guard cell outward rectifier K^+^ (GORK) (Ivashikina et al., 2001; Reintanz et al., 2002) and Stelar K^+^ outward rectifier (SKOR) in *M. trunculata* (Wang et al. 2019) both belonging to the shaker family of potassium channels. A fast-activating outward cation currents (FACC) was also reported from root hairs of *M. truncatula* (Wang et al. 2019). FACC do not seem to depend on shaker family but should be searched in the CNGC or glutamate receptor (GLR) or annexin family of genes (Wang et al. 2019). Two types of anion conductances were described in root hair protoplasts of *M. trunculata* (Wang et al 2019). Slow type anion currents attributed to SLAC channels. These currents resemble the slow type anion currents leading to large depolarization that have been described by dSEVC upon drought stress in *A. thaliana, Phaseolus vulgaris* and *Vigna unguiculata* root hairs (Dauphin et al. 2001). The second type of anion currents described in root hair protoplasts of *M. trunculata*, correspond to anion currents activated upon hyperpolarization. These currents could be supported by ALMT genes. It is noteworthy that hyperpolarization-activated anion conductances have also been reported by dSEVC in root hairs of *M. sativa* (Kurkdjian et al., 2000) and *A. thaliana* (Bouizgarne et al. 2006) but they display no or slow inactivation of currents. This instantaneous activated anionic conductance could serve as an electrical shunt for the H^+^-ATPase which extrudes H^+^ ions into the apoplast and more generally could contribute to short-term electrical and/or calcium signaling, to the regulation of membrane potential and pH gradient across the plasma membrane. Effectively, beside these ion channel activities, the activity of H^+^-ATPases was also described in root hairs of *M. sativa*, date palm and *A. thaliana* by using dSEVC (Kurkdjian et al. 2000, Bouizgarne et al. 2004; 2006). This activity could be supported by autoinhibited plasma membrane H^+^-ATPase genes (AHA). In *A. thaliana* root hairs AHA2 could drive root cell expansion during growth and AHA7 is present in the tip of growing root hair (Hoffman et al. 2019) where they play an important role in root hair formation by mediating H^+^ efflux in the root hair zone (Yuan et al. 2017).

Despite recent efforts, ion conductances in plant root hairs are still poorly studied compared with those in guard cells, for example, particularly in situ. In this study, using dSEVC in *A. thaliana* intact root hairs, we described, the putative role of an outward current presenting a quite chaotic activation upon large depolarization whose physiological role has not been investigated so far. We show that this conductance could serve to hyperpolarize root hairs and switch the whole conductance from a “K^+^ state” dominated by inward rectifying K^+^ currents to a hyperpolarized state due to activation of H^+^-ATPases and anion currents.

## Material and methods

### Plant material

The study was performed on *Arabidopsis thaliana* (Columbia ecotype) seedlings. To allow the germination of the seedlings, after surface sterilization, the seeds were placed in petri dishes on solid media (BS media composed of 5 mM MES, 1 mM CaCl_2_ and 1 mM KCl buffered at pH 5.8 with 5 mM Tris, plus 10 g. l^−1^ Bacto agar BS, Difco, Detroit, MI, USA). Incubation was done at 22 ± 2°C and a photon flux density of 40 µmol m^−2^ s^−1^. The seedlings were then directly perfused with BS. All experiments were carried out at room temperature on 4-day-old seedlings. Young growing root hairs were chosen in zone 1 according to the description of Heidstra et al. (1994).

### Electrophysiology

Impalements were performed with borosilicate capillary glass (Clark GC 150F) micropipettes (resistance: 50 MΩ when filled with 300 mM K-acetate and 10 mM KCl) in BS media. As previously described (Bouteau et al. 1999, Kurkdjian et al. 2000, Dauphin et al. 2001), root hairs were voltage-clamped using an Axoclamp 2B amplifier (Axon Instruments, Foster City, CA, USA) which allows discontinuous single electrode voltage clamp (dSEVC) recordings (Finkel and Redman 1985). The opening or the closure of the channels was obtained with protocols of 40 mV steps from -200 to +80 mV, during 1500 ms, with a resting phase of 1500 ms at a holding potential corresponding to V_m_ before clamping. Voltage and current were digitized with a personal computer fitted with a Labmaster TL-1 acquisition board. The electrometer was driven using pCLAMP software (pCLAMP5.5, Axon Instruments). Experiments were performed at 22 ± 2°C. As previously discussed, (Bouteau et al. 1999), if the shape of root hairs could raise the problem of space clamping (Meharg et al. 1994) the choice of small root hairs of about 15 µm in length [1/10 of the length constant for *A. thaliana* (Meharg et al. 1994)] ensured good voltage clamp conditions without the need for cable correction. We systematically checked to ensure that root hairs were correctly clamped by comparing the protocol voltage values with the ones really imposed. Only a small percentage of root hairs failed to display a linear relationship between theoretical and measured potentials. These root hairs were then abandoned. Kinetics are given as single measurements, representative of at least 10 equivalent tests carried out under the same conditions. Whenever possible, data are given as mean ± SD.

## Results and discussion

Successful impalements were routinely obtained in turgid young root hairs of *A. thaliana*. For a total of 42 root hairs investigated the mean value of the resting membrane potential (V_m_), using BS, was around −110 mV, in the range of what was already described for *A. thaliana* root hairs (Dauphin et al. 2001). The membrane voltages, varied between a minimum value of -50 mV and a maximum of -150 mV. The distribution of the recorded membrane voltages was fitted using moving averages allowing to distinguish two populations (Fig 1A). The root hairs of the two different membrane voltage categories could clearly be separated by their whole current characteristics. These two populations present two different membrane transport states, reminiscent of what has been observed in guard cells (Thiel et al. 1992) with a “diffusion state” dominated by equilibrium potential for K^+^ and a “H^+^ pump state” displaying in addition the activity of H^+^-ATPases. The low polarized population present a mean V_m_ around -80 mV (Fig 1 B,E) and display whole currents dominated by time dependent inward rectifying K^+^ currents (Fig. 1C) probably due to expression of *AKT1* and *AtKC1* (Ivashikina et al., 2001; Reintanz et al., 2002, Wang et al. 2019). This low polarized state could correspond to the “diffusion state” and would indicate a cytosolic K^+^ concentration around 30 mM assuming 1 mM of K^+^ in the external bathing medium. The more hyperpolarized population with a mean V_m_ around -130 mV (Fig 1B,E) would correspond to the “H^+^ pump state” and display whole currents that present an instantaneous activation with weak deactivation (Fig. 1D) that was previously described in root hairs from *M. sativa* and *A. thaliana* (Bouteau et al. 1999, Kurkdjian et al. 2000, Bouizgarne et al. 2006) as the sum of inward rectifying K^+^ currents, plus H^+^-ATPase currents and instantaneous activated anion currents. This anionic conductance could act as an electrical shunt for the H^+^-ATPase linked to expression of *AHA2* and *AHA7* (Hoffman et al. 2019, Yuan et al. 2017) which extrudes H^+^ ions into the apoplast (Barbier et al. 1999).

**Figure 1:**
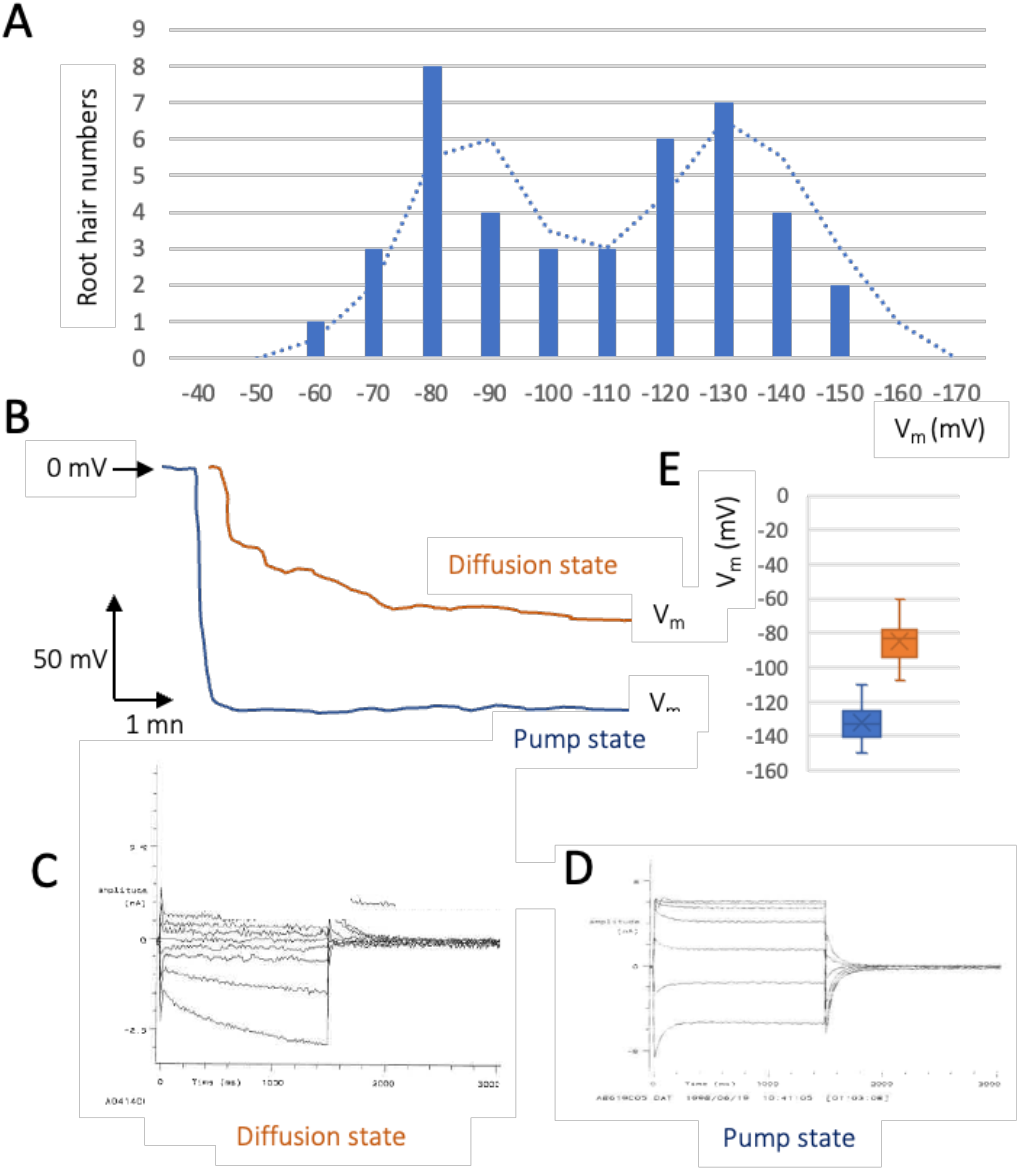
A. Sample and fitted frequency distribution of membrane potentials recorded from a total of 42 root hairs, bathed in BS medium. The recorded voltages were grouped into categories of 10 mV. The theoretical distributions were fitted to the data using moving averages highlighting two populations, a depolarized one and a more hyperpolarized. B. Free running potentials of a root hair of the depolarized population displaying a “diffusion state” dominated by inward rectifying K^+^ currents (C) and free running potential of a root hair of the hyperpolarized population displaying a “pump state” dominated by a more complex whole current (D). E. mean voltages values of the two populations of root hairs.

When considering the less polarized root hairs, ie. the diffusion state dominated by inward rectifying K^+^ currents (Fig 1C), we could frequently observe at positive voltage steps the chaotic and spiky » activation of an outward current (Fig 2A-D). The voltage steps and timing of these current activations are highly variable. The activation could be very fast (Fig 2A,D), resembling the one of the Fast Activating Cation Channels (FACC) described in root hair protoplasts of *M. sativa* (Wang et al. 2019), or appeared only after a few hundreds of milliseconds after the voltage imposition (Fig 2B,C). These outward directed currents present a strong rectification appearing only for the more positive voltage step (+80 mV) (Fig 2 A-C). They could sometimes appear at less depolarized steps, but always positive (Fig 2D) and are systematically irregular and spiky upon voltage imposition, resembling one more time to the FACC described in root hair protoplasts of *M. sativa* (Wang et al. 2019). These characteristics are clearly different from those of the time dependent SKOR currents also described in root hairs (Bouteau et al. 1999, Wang et al. 2019).

**Figure 2:**
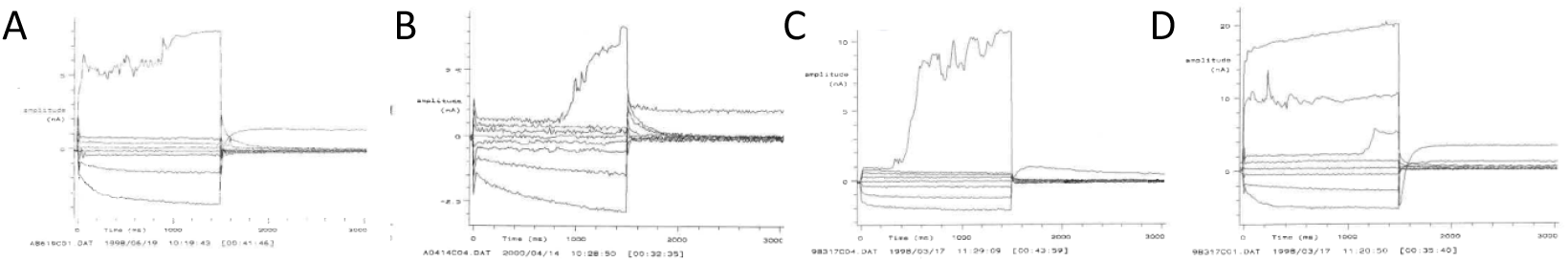
A-D Examples of whole cell currents recorded in the “diffusion state”, displaying typical inward rectifying K^+^ currents and for the +80 mV voltage step (except in E, for 0, +40 and +80 voltage steps) a large outward directed current showing spiky and chaotic activation depending on the root hairs. Note that in B and E this outward directed current could show a fast activation.

The characteristics of this outward current at positive voltage are effectively reminiscent of those described for the FACC of *M. trunculata* root hair protoplasts (Wang 2015, Wang et al. 2019). The latter show spiky activity although their activation is not as chaotic as what we observed. The discrepancy could be due to the *in vivo* current recording conditions (dSEVC) compared with *in vitro* conditions (controlled internal and external conditions in the whole-cell patch clamp). Thus, in this study we chose to use the dSEVC, which allows to work with intact root hairs from seedlings. Indeed, maintaining cell integrity could be of prime importance for the regulation of membrane transport (McCulloch et al. 1997, Thion et al. 1996). Overall, these data lead us to believe that the outward directed spiky current observed in *A. thaliana* root hairs correspond to the FACC described in *M. sativa*, and that it could therefore be supported by cationic currents.

More interestingly, our approaches using dSEVC on intact root hairs also allow us to show that upon activation of this outward spiky current, 95 % of the root hair membrane potentials are hyperpolarized upon the arrest of voltage clamping (green arrow in Fig 3A). During such recordings we also could systematically observe that the current recorded during the resting phase with a holding potential at the V_m_ value recorded before the voltage imposition, is shift from 0 to positive values (red arrow in Fig 3B1) suggesting a modification of the resting value conditions. After the activation of this spiky outward current and the V_m_ hyperpolarization most of the root hairs present a stable hyperpolarized V_m_ and present whole currents (Fig 3B2) resembling the whole currents recorded in the “H^+^ pump state” (Fig 1D) suggesting the activation of H^+^-ATPases and anion channels. In accordance with this change in whole cell currents, the mean V_m_ shift for recording made on the 22 root hairs in diffusion state is of -40 ± 4 mV from -77 mV to -117 mV (Fig 3C).

**Figure 3:**
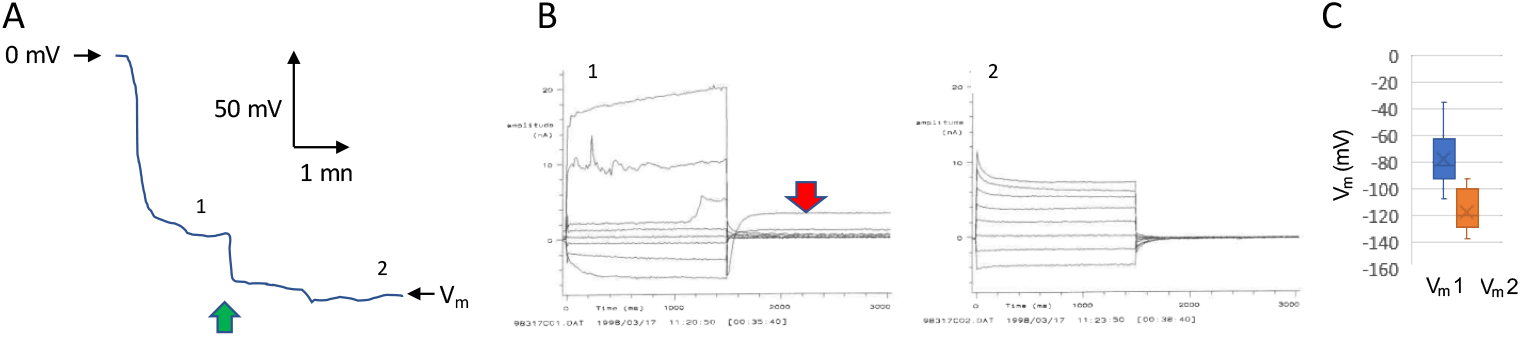
A. Free running potential of a root hair of the depolarized population displaying a “diffusion state” shifting to hyperpolarized state after the protocol of voltage clamping (green arrow). B. Examples of the whole cell currents displaying inward rectifying K^+^ currents plus a large outward directed current showing spiky and chaotic activation at positive voltage steps (B1) and those display after the voltage clamping protocol (B2), thus after the shift to a more hyperpolarized state. Note that the current recorded during the resting phase with a holding potential at the V_m_ value is shift from 0 to positive values (red arrow in B1). C. mean voltage values of the free running potentials recorded for the root hairs before (V_m_1) and after (V_m_2) the voltage clamping protocol.

In 1 on 22 of the root hairs tested the activation of the FACC-like outward current leads only to transient membrane hyperpolarization (Fig 4A, green arrows). In this experiment the whole current is not shift from inward rectifying K^+^ currents to the whole currents recorded in the “H^+^ pump state” (as described in Fig 3) but remains as inward rectifying K^+^ currents (Fig 4B1-3). This explains the only transient hyperpolarization and thus the maintenance of the root hair in a “diffusion state”. This clearly confirms the role of the outward directed spiky current in hyperpolarizing the root hairs but indicates that it is not sufficient to maintain the hyperpolarization, the activation of the H^+^-ATPases and the anion channels being required to maintain the hyperpolarized state.

**Figure 4:**
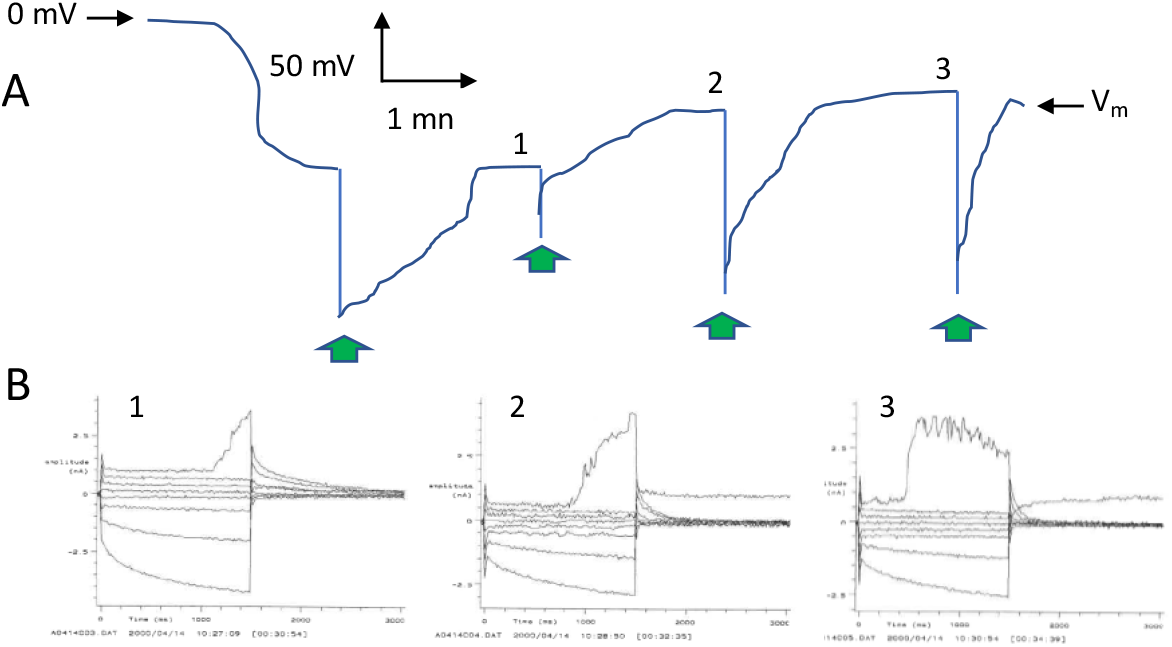
A. Free running potential of a root hair of the depolarized population displaying a “diffusion state” shifting only transiently to hyperpolarized state after the protocol of voltage clamping (green arrow). B. Whole cell currents displaying inward rectifying K^+^ currents plus a large outward directed current showing spiky and chaotic activation at the +80 mV voltage step recorded consecutively (1 to 3 in panels A and B) for this root hair.

Interestingly, we were able to test specific pharmacological tools on the outwardly directed current in a root hair that remained in a “diffusion state” despite the activation of the outwardly directed spiky conductance. It is noteworthy that this outward spiky current is not sensitive to 5 mM cesium nor 5 mM TEA-Cl (Fig 5A,B) which are classic inhibitors of shaker-like K^+^ currents in both plant and animal cells, whereas these molecules efficiently reduce the inward rectifying K^+^ current (Fig 5A). This lack of sensitivity to classical shaker-like K^+^ currents is in accordance with the data obtained in *M. trunculata* root hairs suggesting that the molecular nature of the FACC do not belong to shaker family but should be searched in the CNGC, GLR or annexin families (Wang et al. 2019).

**Figure 5:**
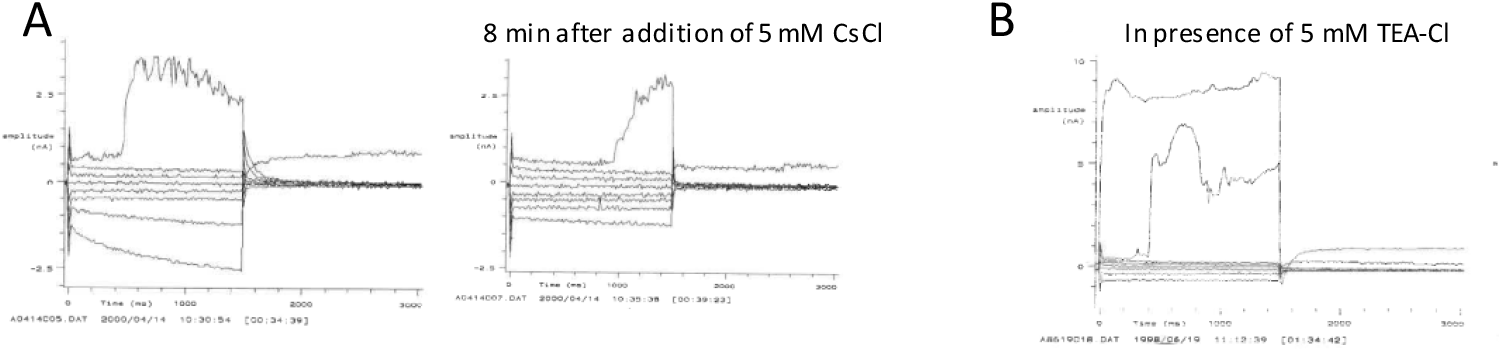
A. Whole cell currents displaying inward rectifying K^+^ currents plus a large outward directed current showing spiky and chaotic activation at the +80 positive voltage step recorded before and 8 min after addition of 5 mM CsCl. B. Whole cell currents recorded for a root hair impaled in BS plus 5 mM TEACl displaying a large outward directed current showing spiky and chaotic activation at the +40 and +80 positive voltage steps.

In conclusion, the role of the FACC is probably to hyperpolarize the root hair during a strong depolarization (in this case artificial). However, the transition from a depolarized state depending only on inward K^+^ currents to a stable hyperpolarized state depending on the activation of H^+^-ATPases and anion channels certainly requires a coupling mechanism between FACC activation and the activation of H^+^-ATPases and anion channels. In the future, a better understanding of the mechanisms at the origin of these interactions will make it possible to establish the key elements of FACC regulation and hence to control cellular responses to environmental stimuli.

